# Generating cell-level protein-expression labels on HE from serial-section immunohistochemistry, with ground-truth-free quality control

**DOI:** 10.64898/2026.06.18.733058

**Authors:** Eunji Jang, Yong-Min Huh

**Affiliations:** YUHS-KRIBB Medical Convergence Research Institute, College of Medicine, Yonsei University, Seoul 03722, Republic of Korea; Department of Radiology, College of Medicine, Yonsei University, Seoul 03722, Republic of Korea

**Keywords:** Computational pathology, Whole slide image registration, Cell matching, Serial-section immunohistochemistry, Label transfer, Matching quality assessment, Protein expression

## Abstract

A laboratory that stains immunohistochemistry (IHC) on a section adjacent to a hematoxylin and eosin (HE) section already holds the material to supervise HE models on its own cases. It goes unused because adjacent sections sample non-identical cells, and residual registration error prevents assigning IHC labels to individual HE cells. We present CellDF (Cell Displacement Field), which turns registered serial-section data into pairs of HE cells and the protein-expression labels measured on the adjacent section. CellDF estimates a locally adaptive residual displacement field through iterated kernel regression over each HE cell’s *K* nearest IHC candidates; a sparse-kernel variant keeps it tractable at whole-slide cell counts, where pairwise matchers are not. The within-tile distribution of these displacements yields two ground-truth-free statistics, the directional scatter *σ*_θ_ and the between-tile angular deviation |*Δθ*|, that localize matching quality more finely than landmark-based target registration error and drive a two-stage filter that withholds labels where matching is unreliable. On 54 same-section HyReCo pairs, *σ*_θ_ correlates only moderately with landmark error and flags localized restaining damage that global error misses; on 30 four-marker Acrobat serial-section cases, the same statistic identifies which IHC marker, if any, lies close enough to HE for cell-level transfer. As a proof of concept, transferred labels trained a cell classifier on HE embeddings that generalized to held-out cells within the sample (F1 0.85, AUROC 0.88). A laboratory can thereby generate cell-level protein-expression labels from its own sections and tune HE-only models on them, with each label set’s reliability read from the data.

## 1. Introduction

Tissue-based diagnosis routinely draws on multiple staining techniques. Hematoxylin and eosin (HE) staining reveals tissue morphology and remains the foundation of diagnostic pathology, while immunohistochemistry (IHC) reports on the spatial expression of specific proteins. In clinical practice, breast cancer assessment combines HE with multiple IHC markers, including estrogen receptor (ER), progesterone receptor (PGR) (Allison et al., 2020), HER2 (Wolff et al., 2018), and the proliferation marker Ki-67 (Nielsen et al., 2021). Automatically transferring IHC marker labels onto individual HE cells would yield per-cell records linking morphology to molecular phenotype across whole slide images (WSI), the pairs of HE cells and their IHC labels that downstream analyses and models depend on. The material for such records is already present wherever IHC is performed routinely, since the two sections are cut and stained in the same laboratory that would use the labels. Yet reliable production of such labels is an open challenge.

Three approaches have been pursued to bridge HE and IHC. Serial-section IHC is the most widely available source of paired HE-IHC data, owing to its routine use in clinical and archival pathology. However, the paired slides are derived from adjacent rather than identical tissue sections. Consequently, differences in cellular composition, together with residual registration error, preclude direct assignment of cell-level labels across slides. Two alternatives attempt to bypass this limitation, each with important trade-offs. Same-section restaining places HE and IHC on the same physical section through destain-restain protocols (Lotz et al., 2023; Remark et al., 2016; Akturk et al., 2020), preserving cell-level correspondence and avoiding the section-to-section variation inherent to serial sections. But antigen degradation during processing limits marker compatibility and constrains the number of stains that can be acquired from a single section, making such datasets scarce and difficult to generate at scale. Virtual staining instead trains generative models to infer IHC-like images directly from HE (Isola et al., 2017; Zhu et al., 2017; Liu et al., 2022; Dubey et al., 2024), with recent approaches replacing direct pixel mappings with richer spatial-context representations (Ma et al., 2026). Yet these methods produce synthesized appearances rather than biological measurements, and the lack of cell-level ground-truth IHC labels makes it difficult both to assess training data quality and to validate generated pseudo-labels. As a result, serial sections, which constitute the largest reservoir of paired HE-IHC data, remain underexploited for cell-level supervision because reliable cell correspondence across sections cannot be established.

Turning serial-section data into a usable labeling source faces three coupled difficulties at WSI scale, even when the IHC section is immediately adjacent to HE. First, modern WSI registration achieves alignment at the micrometer scale (Lotz et al., 2023; Wodzinski et al., 2024b), sufficient to overlay tissue features but not to resolve one-to-one cell correspondence: residual displacement places true correspondents in a neighborhood of candidates rather than at the predicted location, requiring a downstream matcher to recover cell-level pairings. Second, existing methods for cell-to-cell matching are predominantly pairwise correspondence approaches (Myronenko and Song, 2010; Jiang et al., 2024) that scale quadratically with cell count, making full one-to-one matching infeasible at the hundreds of thousands of cells per WSI. A tile-wise workaround (Jeyasangar et al., 2024) mitigates the memory cost but introduces parameter sensitivity and boundary-handling difficulties, while subsampling the correspondences to a tractable count (Nasir et al., 2025) matches only a fraction of cells directly and fills the remainder by interpolation. Third, the resulting matches have no cell-level ground-truth reference, so their reliability must be assessed indirectly; sparse landmark-based metrics such as target registration error (Fitzpatrick et al., 1998) summarize global alignment but cannot localize where matching is trustworthy within a slide. Using the matched cells as a labeling source therefore requires a quality readout that is ground-truth-free and locally resolved.

We address these difficulties with CellDF (Cell Displacement Field), an approach that, built on WSI registration, identifies reliably matched regions of serial-section data without ground-truth correspondences and produces pairs of HE cells and their IHC labels within them. CellDF estimates a locally adaptive residual displacement field through iterated kernel regression on each HE cell’s K nearest IHC candidates; a sparse kernel variant brings the approach into WSI-scale tractability. The displacement field is read in two ways: as cell correspondences that drive IHC label transfer onto HE, and as per-tile statistics that quantify local matching reliability, providing a ground-truth-free quality signal that landmark-based metrics cannot deliver. A statistical outlier filter built on these signals removes unreliable matches before they enter the label set. Because the objective is a reliable set of labeled cells rather than a complete cross-slide alignment, withholding labels where the signal flags unreliable matching is a deliberate part of the design rather than a coverage shortfall. We apply CellDF to two complementary datasets with distinct roles. HyReCo (Lotz et al., 2023) is one of the rare same-section benchmarks: its destain-restain protocol places the same physical cells in both modalities and supplies landmark annotations, making it a controlled reference for matching evaluation. Acrobat (Weitz et al., 2024), with serial-section IHC across four markers per case, represents the practical target setting that motivates this work. Label transfer is demonstrated on an Acrobat sample-marker pair as a proof of concept that the resulting pairs of HE cells and their IHC labels are consistent enough to support downstream learning.

This work treats serial-section IHC as material that a laboratory can turn into cell-level labels on its own cases, which requires both solving cell-level matching and establishing, without ground truth, where the resulting labels can be trusted. The contributions are fourfold. We propose per-tile statistics of a residual displacement field that quantify matching quality with spatial granularity beyond what landmark-based metrics reach and that require no ground-truth correspondences, so the reliability of a label set is read from the data that produced it. Building on these statistics, we add a quality control filter that removes outlier tiles and individual matches before label transfer. To make both of these possible at whole-slide scale, we introduce a non-parametric cell matcher that scales sub-quadratically with cell count and remains tractable at WSI populations where pairwise methods are not. In multi-marker serial-section settings, where the physical adjacency of IHC sections to HE is unrecorded, the same quality metric flags which marker, if any, is close enough to support cell-level transfer. The resulting pairs of HE cells and their IHC labels are consistent enough to train a classifier that generalizes to held-out cells of the same sample, so an HE model can be tuned on labels the laboratory generated from its own material.

## 2. Related Works

### 2.1 Approaches to Bridging HE and IHC at the Cellular Level

Three approaches have been used to combine HE morphology with IHC expression at fine spatial resolution: serial-section IHC, same-section restaining, and virtual staining. Serial-section IHC, in which physically adjacent slices are independently stained with HE and one or more IHC markers, is the standard archival format in clinical pathology and the form represented in datasets such as ACROBAT (Weitz et al., 2024). Because the IHC sections are physically distinct from the HE section, the modalities carry non-identical cell populations in addition to any registration error between them, and cell-level analysis depends on whole-slide registration to bring corresponding tissue regions into approximate alignment. Cell-level correspondence on serial sections has received less attention than registration itself, with recent efforts treating detected nuclei centroids as point sets and applying Coherent Point Drift, refined by graph matching or combined with a shape-aware alignment stage, to recover cell correspondence on top of whole-slide registration (Jiang et al., 2024; Nasir et al., 2025). These methods address the same underlying problem as the cell matcher proposed in this work but rely on pairwise point-set algorithms whose memory cost grows with the product of the two cell populations.

Two workarounds attempt to bypass the misalignment inherent to serial sectioning, each with a cost that limits its use as a scalable labeling source. Same-section destain-restain protocols stain a single physical section with HE, remove the chromogen, and restain with IHC, placing both modalities on the same physical cells. The HyReCo dataset (Lotz et al., 2023) established this approach as a benchmark resource, providing a re-stained subset of 54 section pairs with landmark annotations for evaluating registration. The same destain-restain principle has been extended to four or five sequential IHC markers in protocols such as MICSSS (Remark et al., 2016; Akturk et al., 2020). While these protocols eliminate section-to-section registration uncertainty, the destain-restain cycle degrades antigenicity, the number of markers is bounded by the number of viable cycles, and the specialized chemistry and scanning workflows place these datasets outside routine clinical acquisition. Same-section data is therefore valuable as a controlled benchmark for cross-modal matching algorithms but does not scale as a clinical data source for autolabeling.

Virtual staining circumvents the registration problem entirely by synthesizing IHC-like images from HE alone, using conditional generative models trained on paired or unpaired data. Foundational architectures include pix2pix (Isola et al., 2017) and CycleGAN (Zhu et al., 2017), with later work specializing to breast cancer markers such as HER2 through paired benchmarks and pyramid-pix2pix (Liu et al., 2022) and to multi-marker generation through text-conditioned diffusion (Dubey et al., 2024). Recent work has begun to address misalignment artifacts in paired training data through cascaded registration mechanisms (Ma et al., 2026), and the field has been comprehensively reviewed (Latonen et al., 2024). The generated image is a learned surrogate rather than an actual molecular measurement, and given the scarcity of cell-level real-IHC labels paired with HE morphology, neither training data quality nor the reliability of generated pseudo-labels can be evaluated at the cell level. Published validation therefore remains at the image-quality or slide-level scoring level rather than at cell-by-cell IHC positivity (Latonen et al., 2024).

Exploiting serial-section data for cell-level labeling therefore depends on two capabilities that such workflows still lack: cross-modal cell matching that stays reliable at whole-slide scale, and a means of assessing that reliability without ground-truth correspondences.

### 2.2 Whole-Slide Image Registration for Cross-Modal Alignment

Bringing HE and IHC into the same coordinate system is the prerequisite for any cell-level cross-modal analysis. The field has progressed from classical variational and feature-based pipelines to deep learning approaches, with both lineages cataloged in a recent methodological review (Elhaminia et al., 2025). Classical methods combined affine initialization with patch-based B-spline or elastic deformation (Lotz et al., 2016); despite the subsequent rise of deep learning, intensity-based multi-resolution pipelines remained competitive on the ANHIR challenge (Borovec et al., 2020), which provides the standard cross-stain serial-section benchmark.

Deep learning approaches have improved registration accuracy by learning stain-invariant features and directly estimating dense deformation fields, with VoxelMorph (Balakrishnan et al., 2019) establishing the general unsupervised formulation based on convolutional neural networks (CNNs) in medical imaging. For HE–IHC specifically, the ACROBAT challenge series (Weitz et al., 2024) provides the largest public benchmark, and the winning entry of its 2023 edition combines deep feature matching with intensity-based non-rigid refinement (Wodzinski et al., 2024b); the same framework is distributed as the open-source DeeperHistReg package (Wodzinski et al., 2024a) used in this study.

Even at the micrometer-scale accuracy of state-of-the-art registration, however, the alignment is sufficient to overlay tissue features but not to resolve one-to-one cell correspondences. Landmark-based registration metrics quantify alignment quality at manually selected anchor points but do not characterize residual displacement at the dense, unselected cell positions where one-to-one assignment must be made. Each detected cell sits in a small neighborhood of candidates from the other modality, and the choice among them must be made by a separate matching step. A concurrent line of work has begun to address this directly through nuclei-centroid point-set registration on top of an initial whole-slide alignment, evaluated on the same HyReCo restained data used here (Jeyasangar et al., 2024; Nasir et al., 2025).

### 2.3 Cross-Modal Cell Detection

Cell-level analysis requires detectors that produce consistent results across staining modalities, a non-trivial requirement because most histology cell-detection models are trained on HE alone. Hover-Net (Graham et al., 2019) established CNN-based joint segmentation and classification of nuclei in HE; StarDist (Schmidt et al., 2018) approached the same problem with star-convex polygon representations particularly suited to crowded nuclei. Both are typically trained on HE-only datasets such as PanNuke (Gamper et al., 2019), which provides 189,000 annotated nuclei across nineteen tissue types but no IHC examples. Performance of HE-trained detectors on chromogenic IHC is therefore not guaranteed and is rarely benchmarked in the original methods papers.

A separate strand of work addresses the staining itself rather than the detector, using the Hematoxylin-Eosin-DAB (HED) color deconvolution transform (Ruifrok and Johnston, 2001) to separate hematoxylin counterstain from 3,3’-diaminobenzidine (DAB) chromogen. Computational DAB destaining converts an IHC image into an approximately HE-only image, offering a route to applying HE-trained detectors on both modalities without retraining.

The emergence of large vision foundation models has shifted the cell-detection problem toward a single backbone that can be specialized across stains. The Segment Anything Model (SAM) (Kirillov et al., 2023), pretrained on a billion-mask natural-image dataset, provides a transferable ViT-H (huge Vision Transformer) encoder; CellViT (Hörst et al., 2024) adapts this encoder to nuclear instance segmentation by pairing it with a Hover-Net-style decoder, and CellViT++ (Hörst et al., 2025) extends the approach for energy-efficient and cross-domain deployment. This work uses CellViT++ on both HE and computationally destained IHC. Although the SAM-derived backbone offers stronger cross-domain generalization than HE-only specialists, the model is still trained on HE annotations, and no published benchmark directly evaluates its consistency on paired HE–IHC cells of the same physical section.

### 2.4 Point Set Matching at Scale

Given two sets of detected cells, recovering correspondences between them is a point-set matching problem with a long history in computer vision and a recently re-energized line of work in computational biology. Among algorithms native to point-set registration, Coherent Point Drift (CPD) (Myronenko and Song, 2010) established a widely used non-rigid framework, casting one point set as a Gaussian mixture and driving the other toward alignment through an expectation-maximization procedure, with the deformable variant regularizing the displacement field by motion coherence.

This framework has recently been applied to cell-level cross-modal matching in histopathology, although each uses the resulting correspondences to drive image registration rather than as the final output. Jiang et al. (2024) aligned HE and multiplex immunofluorescence images on ovarian cancer tissue microarrays (TMAs) by establishing one-to-one cell correspondences through graph matching and locality-preserving filtering, then fitting a global affine transform from them, at the cell counts of a TMA core (hundreds to a few thousand cells). A coarse-to-fine engine (Nasir et al., 2025) estimates a global shape-aware rigid alignment, applies CPD for non-rigid refinement over a few thousand matched cells, and interpolates the result into a dense whole-slide deformation field; its accuracy is reported as landmark registration error rather than verified one-to-one cell correspondence. Neither delivers a per-cell correspondence map across the full whole-slide population: the former operates at tissue-microarray scale, and the latter outputs a registration field obtained by interpolation.

Both rely on the same CPD core, whose two iterative steps scale poorly with cell count. The E-step computes a correspondence weight between every target and every source cell, with memory growing as the product of the two populations (*O*(*NM*)); the M-step updates the displacement field by solving an *M* × *M* linear system, since the per-cell weights are coupled through the smoothness regularization, with time growing as the cube of the source count (*O*(*M*^3^)). At whole-slide populations of several hundred thousand cells per modality both become prohibitive on standard hardware, which is why CPD-based methods stay at tissue-microarray scale or fall back to subsampling, tiling, and interpolation. Non-parametric Nadaraya-Watson kernel regression (Nadaraya, 1964; Watson, 1964) removes this coupling: the displacement at each query position is a Gaussian-weighted average of nearby observed displacements, computed independently with no joint solve. The cost per iteration then drops to *O*(*NM*) when correspondences are produced pairwise, and to *O*(*NK*) when each target considers only its *K* nearest sources.

### 2.5 Assessing Matching Quality Without Ground-Truth Correspondences

Image registration is conventionally evaluated by Target Registration Error (TRE), the residual displacement between corresponding landmarks after alignment, with the canonical analysis tracing back to point-based registration theory (Fitzpatrick et al., 1998). In digital pathology, the ANHIR (Borovec et al., 2020) and ACROBAT (Weitz et al., 2024) challenges adopt landmark-based TRE (relative TRE in ANHIR, the median of the per-pair ninetieth-percentile distances in ACROBAT) as their primary ranking metric, and state-of-the-art methods continue to be benchmarked exclusively by these landmark statistics (Wodzinski et al., 2024b). Sparse landmarks summarized by a single number cannot localize where alignment is reliable within a slide, cannot be densified to match the density of cell-level analysis, and are subject to inter- and intra-observer variability in their placement.

A separate literature on registration quality control without ground-truth landmarks has developed in medical imaging, recently surveyed in a scoping review that distinguishes image-based from transformation-based and direct from learned approaches (Bierbrier et al., 2022). Per-voxel error prediction has been pursued through regression forests trained on synthetically perturbed alignments (Sokooti et al., 2019), and the determinant of the Jacobian of the deformation field is the conventional dense plausibility check (Liu et al., 2024). Both bodies of work operate on the deformation field produced by registration rather than on cell-level correspondences, and to our knowledge no published work has reported tile-level quality maps for cell matching specifically.

## 3. Methods

### 3.1 Datasets

#### 3.1.1 HyReCo Dataset (Same-Section Positive Control)

We used the HyReCo-Additional dataset (Lotz et al., 2023), which consists of 54 WSIs from tissue sections that were HE stained, destained, and restained with PHH3 (phospho-histone H3, a mitotic marker). This same-section destain-restain protocol eliminates registration uncertainty by ensuring that both HE and IHC modalities capture the exact same tissue section with identical cell populations.

#### 3.1.2 Acrobat Dataset (Serial-Section Real-World Validation)

We used the Acrobat dataset (Weitz et al., 2024), which contains paired IHC-HE WSIs from serial tissue sections. The dataset includes IHC markers for hormone receptors (ER, PGR), proliferation (Ki-67), and HER2. The images are provided at 10× magnification (approximately 1.0 μm/pixel). The Acrobat dataset is split into training, validation, and test sets; only the training set provides multiple IHC markers per HE slide. From this training set, we selected the first 30 samples (sorted by sample name) for which all four IHC markers were available, yielding 120 HE–IHC pairs.

### 3.2 Preprocessing: Image Registration and Cell Detection

#### 3.2.1 Format Standardization

All WSIs were standardized to OpenSlide-compatible formats. Magnification and microns-per-pixel (MPP) values were read from image headers; low-resolution images (≥ 1.0 μm/pixel, such as the Acrobat 10× dataset) were upsampled to approximately 0.5 μm/pixel (equivalent to 20× magnification) by bicubic interpolation.

#### 3.2.2 WSI Registration

We employed DeeperHistReg (Wodzinski et al., 2024a), a deep learning-based registration method, to align the IHC (Source) and HE (Target) WSI pairs. Registration was performed at reduced resolution (initial loading at 1/20 scale, with subsequent processing at 2048×2048 pixels) using the default_initial_nonrigid configuration, which performs rigid initialization followed by iterative non-rigid refinement. The output is a warped Source image transformed into the Target coordinate system.

#### 3.2.3 Cell Detection

We used CellViT++ (Hörst et al., 2025) with the SAM (Kirillov et al., 2023) ViT-H backbone to detect and segment cells in the HE and warped IHC images. The HE image was processed directly, while for the warped IHC image, we first removed the DAB chromogen via HED color deconvolution following the standard scikit-image implementation, followed by contrast enhancement on the H and E channels. For each detected cell, CellViT++ provides centroid coordinates, bounding boxes, segmentation contours, and a 1280-dimensional feature embedding extracted from the SAM ViT-H encoder.

### 3.3 Cell Matching

#### 3.3.1 Matching Strategies

Given Target cells T = {*T*_1_, …, *T*_n_} from HE at positions **x**_i_ and Warped cells G = {*W*_1_, …, *W*_m_} from IHC at positions **y**_j_, we seek one-to-one correspondences to transfer IHC labels from W to T. Pairing a target *T*_i_ with a warped candidate *W*_j_ defines a displacement vector **d** = **y**_j_ − **x**_i_ (warped minus target position). We describe the proposed matcher first, then four baseline matchers compared against it.

CellDF is a scalable non-parametric approach that estimates a *locally adaptive* displacement field rather than a single global translation (Supplementary Fig. 1). For each target cell, the *K* nearest warped candidates within a maximum matching distance are retrieved, yielding up to *Kn* candidate displacement vectors in total. The geometric median of all candidates provides a global initial estimate. Each target cell selects the candidate closest to this consensus. Then, over *L* iterations, the displacement field is re-estimated locally using Nadaraya-Watson kernel regression:

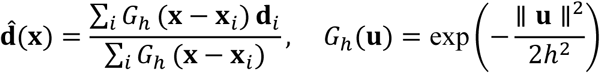

**Supplementary Fig. 1.**
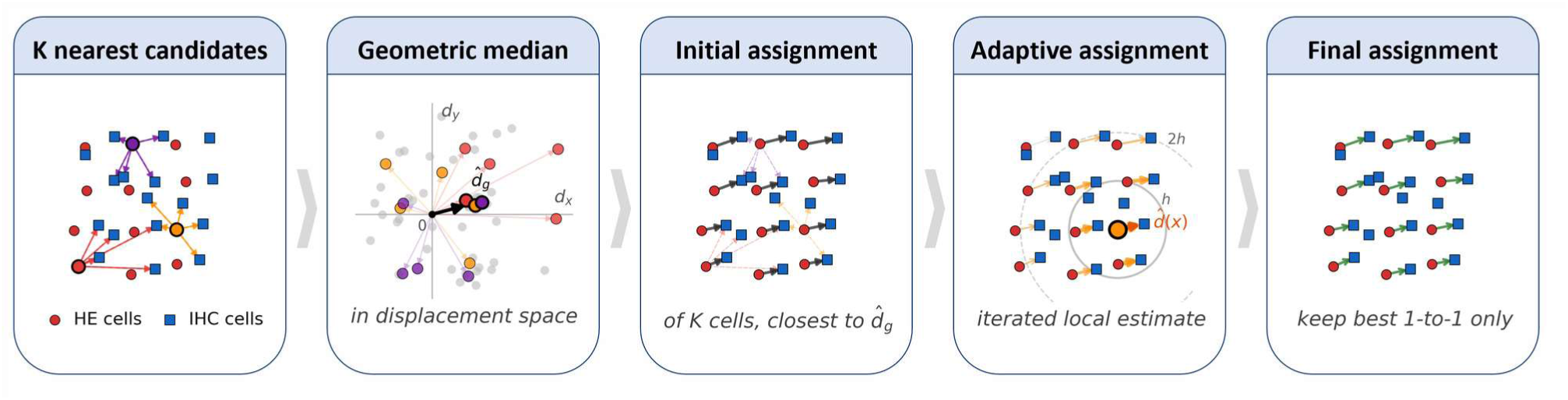
CellDF (Cell Displacement Field) algorithm schematic. CellDF is a scalable, non-parametric cell correspondence method that estimates a locally adaptive displacement field via Nadaraya-Watson kernel regression. Synthetic cells illustrate the five stages: HE (red circles) and IHC (blue squares). **(a)** For three highlighted targets, the *K* nearest IHC cells are retrieved as candidate matches (colored arrows; *K* = 5 shown). **(b)** The *Kn* candidate displacements plotted in displacement space; the geometric median 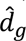 (bold black arrow) provides a robust global initial estimate. Per highlighted target, the candidate closest to 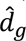 is marked with a thick black border. **(c)** Initial assignment: each target selects its candidate closest to 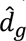; non-selected candidates shown as faint dashed arrows. **(d)** Adaptive step for one focus target (orange border): a Gaussian kernel with bandwidth ℎ (dashed circles at ℎ, 2ℎ) weights neighboring displacements to produce a local estimate 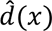 (bold orange arrow). **(e)** After *L* iterations of local re-estimation and greedy 1-to-1 resolution, final matches (green arrows) are retained.

where 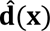 is the locally estimated displacement at a query target position **x**, the sum runs over the current matches, **d**_i_ is the displacement of matched target *i* at position **x**_i_, and *G*_h_ is the isotropic Gaussian kernel of bandwidth ℎ. Each target cell re-selects the candidate best agreeing with the updated local prediction. Remaining one-to-one conflicts are resolved by retaining the match with smallest deviation from the local field. This dense kernel regression computes pairwise distances among all matched positions, which is feasible at patch scale but prohibitive at WSI scale; for WSI-level application, we use a sparse variant that truncates the kernel at radius 3ℎ, evaluating only spatially nearby points. We used *K* = 5, *L* = 3, ℎ = 50 *μ*m for HyReCo and 100 *μ*m for Acrobat, and a maximum candidate matching distance of 25 *μ*m for HyReCo and 50 *μ*m for Acrobat. Convergence in *L* and sensitivity to ℎ on the 54-sample HyReCo cohort are reported in Supplementary Fig. 2.

**Supplementary Fig. 2.**
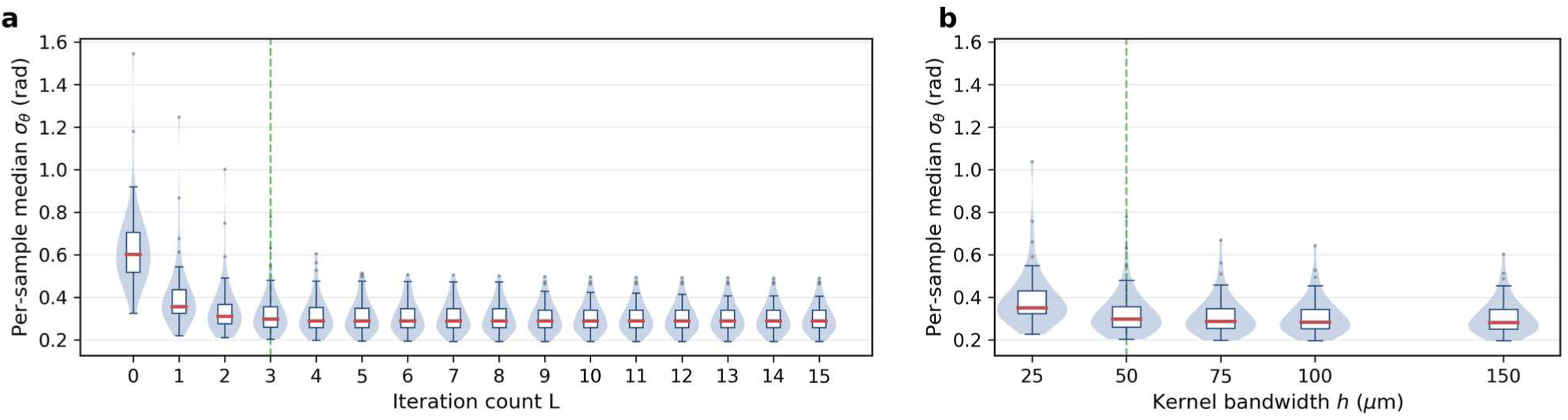
CellDF sensitivity to refinement iterations *L* and kernel bandwidth ℎ across 54 HyReCo samples. Each violin, with overlaid box and median, shows the distribution across all 54 HyReCo HE–IHC pairs of the per-sample median tile-level *σ*_θ_ at one parameter value. **(a)** As a function of the number of CellDF refinement iterations *L* ∈ {0,1, …, 15}. **(b)** As a function of the Nadaraya–Watson kernel bandwidth ℎ ∈ {25,50,75,100,150} *μ*m, with *L* = 3 held fixed. Green dashed lines mark the HyReCo defaults used elsewhere in the paper (*L* = 3, ℎ = 50 *μ*m).

The four baselines span the principal design choices a residual-displacement matcher can make. First, nearest-neighbor matching imposes no spatial coherence, pairing each target with its Euclidean-nearest warped centroid and each warped with its nearest target and retaining only mutual best matches; the queries are accelerated by a KDTree (Bentley, 1975).

Second, CPD (Myronenko and Song, 2010) is a parametric deformable point-set registration method that models one point set as a Gaussian mixture and drives the other toward alignment through a globally coupled, Gaussian-regularized displacement field. We used the deformable variant with hyperparameters tuned on pilot data (*α* = 0.01, *β* = 100, *σ*_0_ = 30 px, *w* = 0), with mean-centering applied for numerical stability.

Third, optimal transport forms correspondences by computing a coupling between the two point sets. We use a coherence-aware transport cost: at each of *L* = 3 outer iterations the cost compares each warped cell against the predicted position of each target under the displacement field estimated from the previous round’s matches, 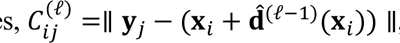, with 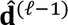 the Nadaraya-Watson kernel-smoothed displacement field (plain Euclidean distance at the first iteration); a mutual-best argmax of the resulting coupling yields one-to-one matches. Because the two cell populations differ in size (*n* ≠ *m*) and need not match completely, the coupling is computed with an Unbalanced Optimal Transport formulation (Chizat et al., 2018), solved by the majorization-minimization (MM) solver of the POT library (Flamary et al., 2021) (*ε* = 0, no entropic regularization; *λ* = 10, marginal relaxation strength); we denote this baseline UOT-MM.

Finally, Random Sample Consensus (RANSAC) (Fischler and Bolles, 1981) fits a single global translation rather than a spatially-varying field. From the *Kn* candidate pairs formed by each target’s *K* nearest warped candidates, random 3-pair subsets estimate a consensus translation (mean displacement) over *T* iterations, retaining the model with the most inliers (deviation < 30 pixels) and taking the inlier mean as the refined consensus. Each target then selects the candidate closest to this consensus, with one-to-one conflicts resolved by deviation and no local re-estimation following. We used *K* = 5 and *T* = 500.

#### 3.3.2 Computational Scaling

The five methods differ substantially in computational scaling, summarized in Table 1. Let *n* and *m* denote the number of target and warped cells, respectively.

**Table 1.** Computational complexity of cell matching methods. Time includes both candidate search (where applicable, the (*n* + *m*)log*m* term from KDTree build and *K*-NN queries) and the core algorithm. Memory is peak space required during execution, independent of time. *n*: target cells, *m*: warped cells, *L*: outer coherent-cost iterations for UOT-MM and refinement iterations for CellDF, *L*_mm_: UOT-MM inner solver iterations per outer round, *L*_em_: CPD EM iterations, *T*: RANSAC iterations, *K*: nearest neighbors. CellDF (dense) is used at patch scale; CellDF (sparse) truncates the kernel at 3ℎ for WSI-scale application.

| Method | Time | Memory |
| --- | --- | --- |
| Nearest (KDTree) | $O((n + m)\log m)$ | $O(n + m)$ |
| CPD | $O(L_{em} \cdot m^3)$ | $O(nm + m^2)$ |
| UOT-MM | $O(L \cdot L_{mm} \cdot nm)$ | $O(nm)$ |
| RANSAC | $O((n + m)\log m + Tn)$ | $O(n + m)$ |
| CellDF (dense; patch) | $O((n + m)\log m + Ln^2)$ | $O(n^2)$ |
| CellDF (sparse; WSI) | $O((n + m)\log m + Ln\log n)$ | $O(n + m)$ |

Among the five methods, Nearest, RANSAC, and CellDF achieve sub-quadratic scaling by combining *K*-NN candidate search with linear-time core algorithms, whereas UOT-MM and CPD are pairwise in *n* and *m*. UOT-MM and CPD are therefore not generally tractable at WSI-scale populations on standard hardware. Patch-wise application, subdividing the WSI into tiles and running UOT-MM or CPD per tile, could circumvent this but falls outside the scope of the present study.

#### 3.3.3 Per-Tile Displacement Statistics

Rather than constructing ground-truth correspondences for matching evaluation, we assess matching quality through displacement consistency, the spatial smoothness of the resulting displacement field. We partition the WSI into non-overlapping tiles (1024 × 1024 pixels) and, for each tile with at least three matches, summarize the within-tile displacement distribution by its directional statistics: the circular median angle 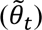, the directional center of the within-tile displacements, and the directional scatter (*σ*_θ_, the circular standard deviation of displacement angles), which measures directional coherence, so that lower values indicate matches sharing a common displacement direction. As a complementary descriptor we also compute the magnitude scatter (*σ*_m_, the standard deviation of displacement magnitudes), capturing the uniformity of displacement distances. WSI-level matching quality is then summarized by statistics of these per-tile values, namely the median and the interquartile range (IQR), together with distributional plots such as violin plots or spatial heatmaps.

### 3.4 Matching Quality Control

At WSI scale, CellDF matching produces hundreds of thousands of correspondences, a fraction of which are unreliable due to local registration failure, tissue folding, or cell or tissue present in only one modality. We apply a two-stage outlier filter based on the Median Absolute Deviation (MAD) to remove these matches before label transfer.

#### 3.4.1 Tile-Level Filtering (Stage 1)

Using the per-tile circular median angle 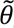_*t*_ and directional scatter *σ*_θ_ defined above, we compute the global displacement direction 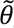_*g*_ as the circular median of all 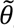_*t*_. In the thresholds below, *σ*_MAD_ denotes the MAD-based robust standard-deviation estimate. Two criteria flag outlier tiles: (A) tiles whose absolute angular deviation |*Δθ_t_*| = |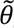_*t*_ − 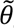_*g*_| exceeds med(|*Δθ*|) + 2 *σ*_MAD_(|*Δθ*|), indicating displacement directions inconsistent with the global field; and (B) tiles whose *σ*_θ,t_ exceeds med(*σ*_θ_) + 2 *σ*_MAD_(*σ*_θ_), indicating internally scattered displacements. Tiles flagged by either criterion are removed entirely. The two criteria are complementary: criterion A captures between-tile deviation from the global displacement field, while criterion B captures within-tile angular scatter.

#### 3.4.2 Match-Level Filtering (Stage 2)

Within each surviving tile, individual matches are filtered by the Euclidean residual *r* between each match’s displacement and the tile’s median displacement vector. Matches whose *r* exceeds med(*r*) + 2 *σ*_MAD_(*r*) are removed, targeting sporadic mismatches that passed tile-level screening. Together, the two stages remove spatially coherent regions of poor registration (Stage 1) and isolated outliers within well-registered regions (Stage 2).

### 3.5 Label Transfer

Because the IHC protocols in the benchmark datasets (HyReCo, Acrobat) use DAB as the chromogen, we applied the HED transform to the original warped Source image to extract the DAB channel and thresholded it to generate a binary DAB mask. For each detected cell in the warped Source, we calculated the overlap ratio between its bounding box and the DAB mask.

Cells are labeled Positive if the overlap ratio is ≥10% and Negative if it is 0%. Cells with overlap ratios between 0% and 10% are excluded from labeling as Ambiguous.

For each matched pair (*T*_i_, *W*_j_) surviving the two-stage quality control, we transfer the IHC label from *W*_j_ to *T*_i_ if the label is Positive or Negative; *T*_i_ is left unlabeled if *W*_j_ is Ambiguous. Target cells without a transferred label constitute the prediction set for label extension: those whose match was removed during quality control, those matched to an Ambiguous cell, and any unmatched cells.

### 3.6 MLP Classification

We train a multilayer perceptron (MLP) on the CellViT++ cell embeddings of labeled target cells to predict IHC labels for the prediction set. For evaluation, we adopt a spatial cell-count split rather than a random partition: the WSI is partitioned horizontally at the y-median of labeled cells into Top and Bottom halves with equal labeled-cell counts, so that adjacent cells with spatially correlated embeddings do not straddle the train and test sets; two MLPs are trained with Top/Bottom reversed as train/test, and performance is reported as F1 score, area under the receiver operating characteristic curve (AUROC), accuracy, precision, and recall on the held-out halves, treating transferred labels as ground truth. For prediction, a separate MLP trained on all labeled cells is applied to the prediction set.

## 4. Results

### 4.1 Preconditions for CellDF: Registration and Cell Detection

CellDF requires two preconditions on the input: (i) residual displacement after WSI-level registration that places the true correspondent within each target cell’s *K*-nearest warped candidates, and (ii) robust cell detection in both modalities so that candidate sets are non-empty.

On the 54-sample HyReCo dataset, the per-sample median TRE after DeeperHistReg registration had a cohort median of 2.5 μm (IQR: 2.1–3.2 μm; 48 of 54 below 8 μm). CellViT++ detected 28.7 million HE and 36.9 million IHC cells (median per sample: 548K and 662K), with Pearson *r* = 0.90 (*p* < 10^−19^) between HE and IHC cell counts. The sub-cellular residual displacement and consistent cross-modal detection together confirm both preconditions.

### 4.2 Patch-Level Cell Matching: Proof of Concept

As a proof of concept for cell-level matching, we selected a representative sample (case 281 in HyReCo; median TRE = 6.5 μm; Fig. 1a–c) and focused on a 1024 × 1024 pixel region (∼256 × 256 μm). Within this region, CellViT++ detected 276 target (HE) and 254 warped (IHC) cells (Fig. 1d–e). Overlaying both sets of contours showed that, despite registration, corresponding cells remained spatially displaced rather than co-located, so cell-level correspondence cannot be assigned by proximity alone and requires an explicit matching step (Fig. 1f).

**Fig. 1.**
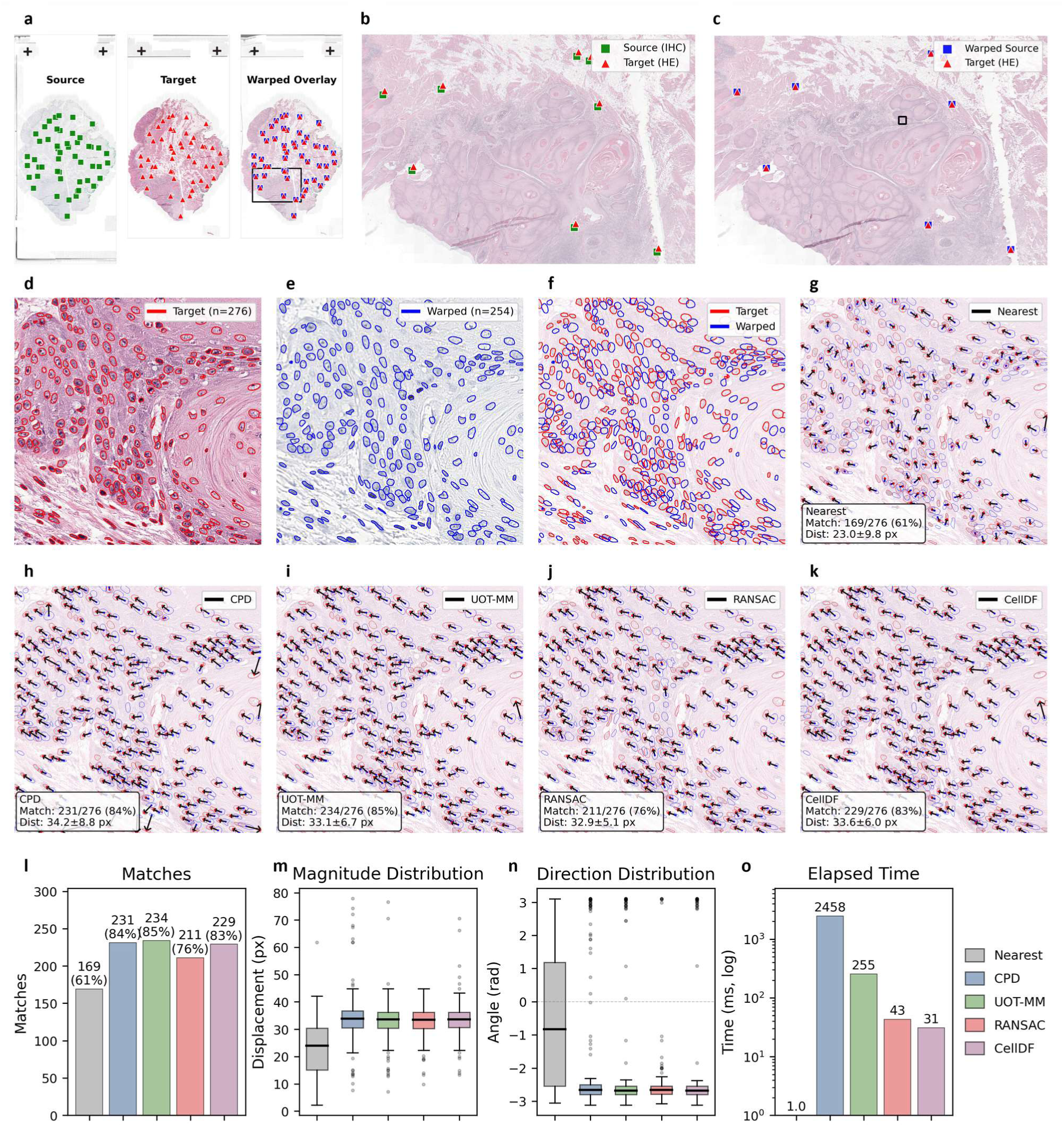
Proof-of-concept for patch-level cell matching. A 1024 × 1024 pixel region from Sample 281 (HyReCo same-section IHC) was used to compare five matching approaches. **(a)** WSI-level overview showing source (IHC), target (HE), and warped overlay with anatomical landmarks; the black box indicates the region enlarged in (b,c). **(b)** Overlay before registration with source (green) and target (red) landmarks. **(c)** After registration; the black box marks the detail region used in (d–k). **(d)** Target patch with cell contours (red, n = 276). **(e)** Warped IHC patch with cell contours (blue, n = 254). **(f)** Contour overlay showing spatial correspondence between target (red) and warped (blue) cell populations on a faded target background. **(g)** Mutual-best-match nearest-neighbor (Nearest, KDTree) result on the same overlay; arrows connect matched pairs. **(h–k)** Matching results for CPD (h), UOT-MM with Gaussian-RBF coherent cost (UOT-MM, i), RANSAC fitting a single global translation (j), and CellDF (k). **(l–o)** Summary metrics across the five methods: match count (l), displacement magnitude distribution (m), angular distribution (n), and elapsed time (o, log scale, milliseconds).

We evaluated five matching strategies on this region (Fig. 1g–k). Only the Nearest (KDTree) baseline produced scattered arrows; CPD, UOT-MM, RANSAC, and CellDF all produced largely coherent vector fields and matched similar numbers of cells (Fig. 1l). The displacement magnitude and angle distributions (Fig. 1m,n) confirmed that Nearest was substantially more dispersed, while CPD, UOT-MM, RANSAC, and CellDF showed narrower, comparable spreads.

Although CPD, UOT-MM, RANSAC, and CellDF achieved similar matching quality at this scale, RANSAC and CellDF ran more than an order of magnitude faster than UOT-MM and nearly two orders of magnitude faster than CPD (Fig. 1o), and the pairwise scaling of CPD and UOT-MM precludes WSI-scale application. At WSI scale, residual registration error varies spatially across the slide; we therefore applied both RANSAC and CellDF to full WSI matching to compare their behavior in this regime.

### 4.3 WSI-Level Matching: RANSAC vs. CellDF

We applied both RANSAC and CellDF to the full WSI of Sample 281, computing per-tile match rate, directional scatter (*σ*_θ_), and magnitude scatter (*σ*_m_) across 6,595 tiles of 1024 × 1024 pixels. We visualized each metric as a spatial heatmap overlaid on the WSI (Fig. 2a,b). RANSAC heatmaps showed relatively modest spatial variation in both scatter metrics, whereas CellDF heatmaps exhibited substantially larger local variation, most prominently in *σ*_θ_. The RANSAC–CellDF agreement map (Fig. 2c) tracked the scatter heatmaps: in low-scatter tiles, the two methods produced similar matching results, while they diverged in tiles of elevated scatter.

**Fig. 2.**
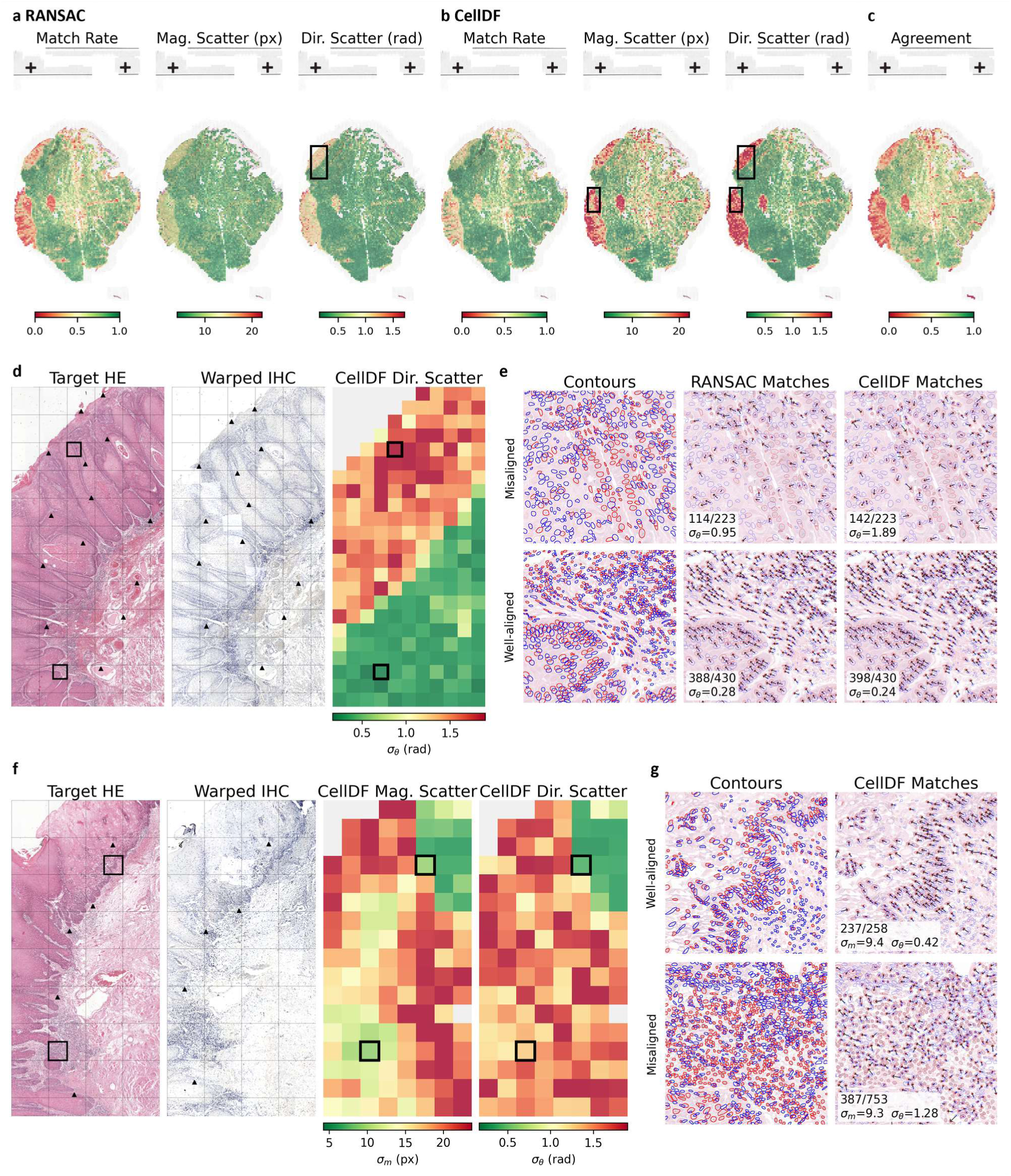
WSI-level tile metric comparison between RANSAC and CellDF. Analyses were performed on Sample 281 (HyReCo same-section IHC) with 1024 × 1024 pixel tiles. **(a)** RANSAC tile-level heatmaps for match rate, magnitude scatter (standard deviation, *σ*_m_), and directional scatter (circular standard deviation, *σ*_θ_). **(b)** CellDF tile-level heatmaps for the same three metrics. **(c)** RANSAC–CellDF agreement map. The boxed region in (a,b) indicates the area enlarged in (d,e); the boxed region in (b) indicates the area enlarged in (f,g). **(d)** Enlarged region: target, warped, and CellDF *σ*_θ_ heatmap. Black triangles indicate manually annotated landmarks. **(e)** Representative patches from (d): well-aligned and misaligned patches with contour overlays and match arrows for RANSAC and CellDF, annotated with per-method *σ*_θ_, match count, and target cell count. **(f)** Second enlarged region: target, warped, CellDF *σ*_m_, and CellDF *σ*_θ_. Black triangles indicate manually annotated landmarks. **(g)** Representative patches from (f): well-aligned and misaligned patches with contour overlays and CellDF match arrows, annotated with *σ*_θ_, *σ*_m_, match count, and target cell count.

To investigate the source of this variation, we enlarged a region containing both low and high CellDF *σ*_θ_ tiles (Fig. 2d,e). Manually placed landmarks on the target (HE) and warped (IHC) images revealed that despite globally successful registration, local alignment quality varied within this region: some tiles showed well-overlapping landmarks while others exhibited visible displacement. Comparing these landmarks with the CellDF *σ*_θ_ heatmap, *σ*_θ_ appeared to track local registration quality: tiles with well-overlapping landmarks had low *σ*_θ_, while tiles with visible landmark displacement had high *σ*_θ_. We therefore examined two representative tiles in detail (Fig. 2e). The bottom tile, where HE and warped IHC cell contours largely overlapped, showed spatially smooth arrow patterns from both methods (RANSAC *σ*_θ_ = 0.28 rad, CellDF *σ*_θ_ = 0.24 rad). The top tile fell in a region of insufficient registration, where the overlaid contours came from largely different cell populations. In this tile, CellDF reached *σ*_θ_ = 1.89 rad while RANSAC stayed at 0.95 rad, with RANSAC’s displacement vectors showing a less disordered pattern. This reflects a property of the single global translation model: constraining every match toward one consensus vector suppresses *σ*_θ_ in locally misaligned regions.

We next examined whether *σ*_θ_ and *σ*_m_ capture distinct information. They diverged in specific regions; we enlarged a region where *σ*_θ_ was high but *σ*_m_ was relatively low (Fig. 2f,g). Manual landmarks confirmed poor local registration in this area. Closer inspection revealed that this area combined poor local registration with high cell density; in such conditions, incorrectly matched pairs still produce small displacement magnitudes, suppressing *σ*_m_ while *σ*_θ_ remains elevated.

These observations suggest that CellDF, applied after WSI-level registration, can reveal local registration quality that global metrics such as median TRE may not capture. Among the methods examined, CellDF reflected local misalignment more reliably than RANSAC, and *σ*_θ_ proved less susceptible to confounding factors than *σ*_m_. To evaluate generalization beyond a single sample, we applied CellDF with *σ*_θ_ to all 54 HyReCo samples and compared per-tile *σ*_θ_ distributions against landmark-based median TRE.

### 4.4 HyReCo: Same-Section Matching

We applied CellDF matching to all 54 HyReCo samples, computing per-tile *σ*_θ_ across 220,273 tiles (1024 × 1024 pixels; tiles with fewer than 3 matched cells were excluded from *σ*_θ_ computation). Figure 3a shows the per-landmark TRE distribution for each sample sorted by median TRE (median 40 landmarks per sample, range 10–100), and Figure 3b shows the corresponding per-tile *σ*_θ_ distributions in the same order. The per-sample median *σ*_θ_ showed a moderate positive correlation with the per-sample median TRE (Spearman *ρ* = 0.38, p = 0.004), indicating that the two metrics capture related but not identical aspects of alignment quality. Most samples exhibited *σ*_θ_ distributions comparable to Sample 281. However, six samples had *σ*_θ_ distributions significantly higher than Sample 281 (Mann-Whitney U, one-sided, p < 0.05; red in Fig. 3b). Among these six elevated-*σ*_θ_ samples, four had low median TRE (2.0–3.2 μm).

**Fig. 3.**
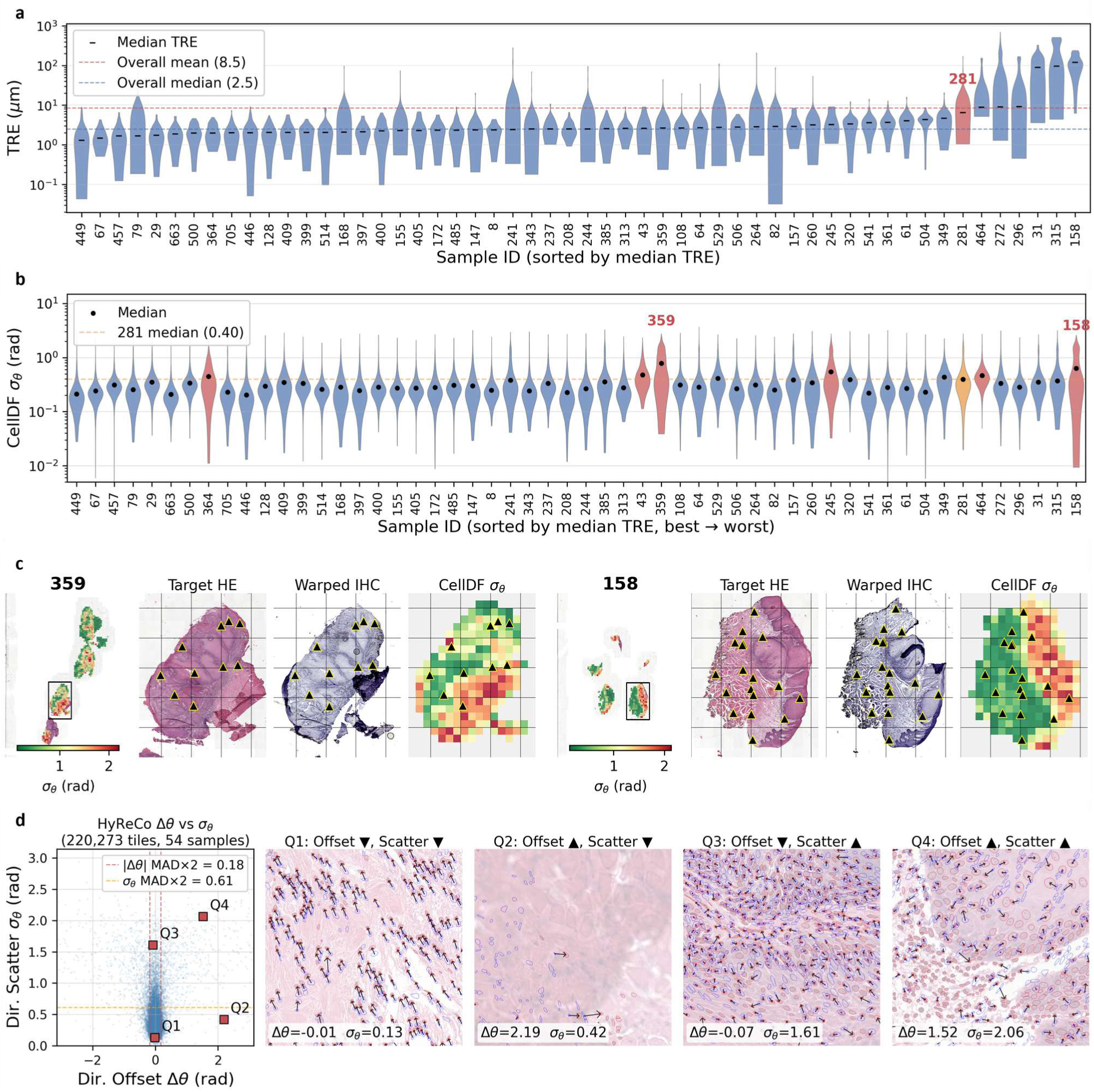
Dataset-wide CellDF *σ*_θ_ analysis across 54 HyReCo samples. **(a)** Per-sample TRE violin plots (log scale), sorted by per-sample median TRE. Each violin shows the per-landmark TRE distribution; horizontal dashes indicate per-sample medians. Sample 281 (red) is the reference sample used in Figures 1–2. Dashed lines: cohort mean and median of the per-sample median TRE. **(b)** Per-sample CellDF *σ*_θ_ violin plots in the same sample order. Blue: not significantly different from Sample 281; orange: Sample 281 (reference); red: significantly higher than Sample 281 (Mann-Whitney U, one-sided, p < 0.05). Dashed line: Sample 281 median *σ*_θ_. **(c)** Two samples with elevated *σ*_θ_ distributions (359, left; 158, right). For each sample: WSI overview with *σ*_θ_ heatmap overlay and region box, target region, warped region, and tile-level *σ*_θ_ heatmap of the boxed region. Black triangles mark landmark positions: HE landmarks on the target region and *σ*_θ_ heatmap, and the displacement-field-warped IHC landmarks on the warped region. **(d)** Left: per-tile directional offset (*Δθ*) versus *σ*_θ_ scatter plot pooled across all 54 samples; *Δθ* is the signed difference between each tile’s median displacement angle and the sample-wide circular median. Dashed lines indicate MAD×2 outlier thresholds for |*Δθ*| (red) and *σ*_θ_ (orange). Labeled squares (Q1–Q4) mark four representative tiles shown at right, corresponding to the four combinations of low/high |*Δθ*| and low/high *σ*_θ_. Right: HE tile patches with CellDF match arrows (Q1, Q4 from Sample 158; Q2, Q3 from Sample 359). Red contours: target cells; blue contours: warped cells; black arrows: CellDF matches.

We examined two samples with the highest *σ*_θ_ (Fig. 3c). Despite differing markedly in median TRE (Sample 359: 2.6 μm; Sample 158: 121 μm), both showed similarly elevated median *σ*_θ_ (0.78 and 0.63 rad respectively) and visible tissue damage in the warped IHC image. The tile-level *σ*_θ_ heatmaps captured both the mismatches at sites of visible tissue damage and the surrounding regions whose registration was degraded as a consequence, flagging Sample 359 despite its globally moderate median TRE. Conversely, even the best-matched samples (those with the lowest per-sample median *σ*_θ_) contained individual tiles with locally elevated *σ*_θ_ despite globally excellent registration (median TRE near 2 μm); direct inspection of the matches in these tiles confirmed local degradation (Supplementary Fig. 3). Taken together, the only moderate *σ*_θ_–median TRE correlation, the localization of tissue damage missed by global median TRE, and the local *σ*_θ_ variation present even in the best-matched samples indicate that *σ*_θ_ resolves matching quality at the tile level, beyond what sparse landmark-based metrics such as median TRE capture.

**Supplementary Fig. 3.**
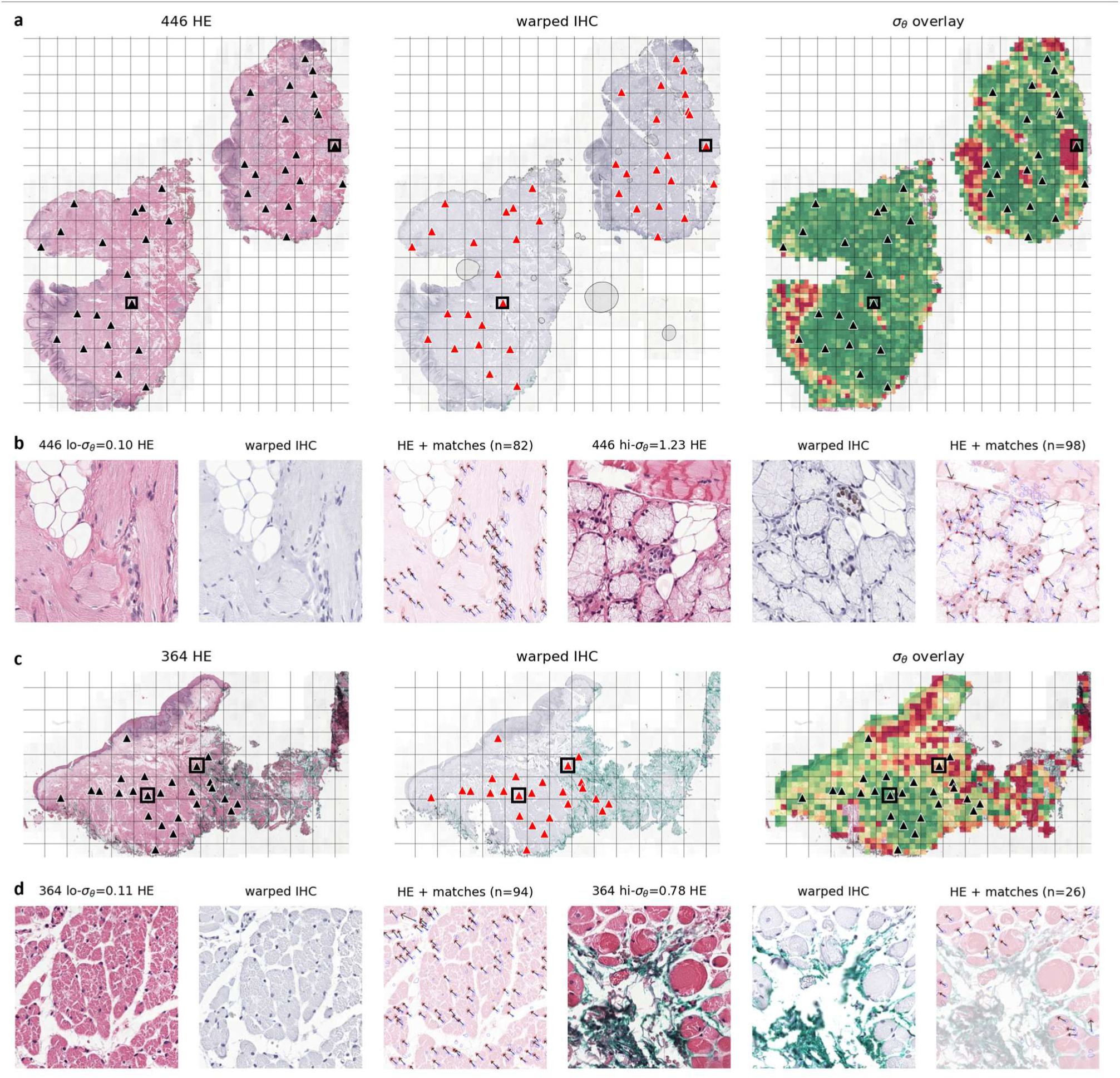
Local *σ*_θ_ variation within the best-matched HyReCo samples. The two HyReCo HE– IHC pairs with the lowest per-sample median *σ*_θ_ (sample 446, a–b; sample 364, c–d; each comprising one to two tissue pieces). **(a, c)** Whole-tissue rows: target HE, displacement-field-warped IHC, and the per-tile *σ*_θ_ heatmap overlaid on HE (shared color scale). A common reference grid is overlaid on all three panels to aid landmark comparison. Black triangles mark HE landmarks (HE and *σ*_θ_ panels); red triangles mark the same landmarks warped through the registration displacement field (warped IHC panel); black boxes locate the landmarks of the two detail tiles. **(b, d)** Detail rows: the two landmark-containing tiles with the lowest (left) and highest (right) *σ*_θ_, each shown as HE, warped IHC, and a match overlay (faded HE with red target-cell contours, blue warped-cell contours, and black CellDF match arrows).

The detailed observations of tissue-damaged slides above motivated complementing *σ*_θ_ with an additional metric. To capture not only within-tile but also between-tile directional consistency, we defined per-tile |*Δθ*|, the absolute deviation of the tile’s median displacement angle from the sample-wide circular median angle, and examined its joint distribution with *σ*_θ_ across HyReCo tiles (Fig. 3d). Perhaps reflecting HyReCo’s same-section nature, the majority of tiles fell in Q1 (low *σ*_θ_ and low |*Δθ*|). Tiles with low *σ*_θ_ but elevated |*Δθ*| were observed at low frequency (Q2). As *σ*_θ_ increased, the range of |*Δθ*| widened correspondingly (Q3, Q4). Zoom-in examples from Q2, Q3, and Q4 tiles (Fig. 3d, right) illustrate representative cases of matching

quality compromised by unmet CellDF preconditions.

### 4.5 Acrobat: Serial-Section Matching

Extending this analysis to serial-section data, we applied the same CellDF pipeline to 30 Acrobat training samples with all four IHC markers available (120 HE–IHC pairs); per-tile *σ*_θ_ distributions shifted substantially toward higher values compared to HyReCo (Fig. 4a). The Acrobat pairs spanned 0.78 to 1.80 rad, whereas the HyReCo reference Sample 281 had a median *σ*_θ_ of 0.4 rad. Compared to HyReCo, both the |*Δθ*| and *σ*_θ_ distributions broadened substantially (Fig. 4b), likely reflecting the nature of serial-section data.

**Fig. 4.**
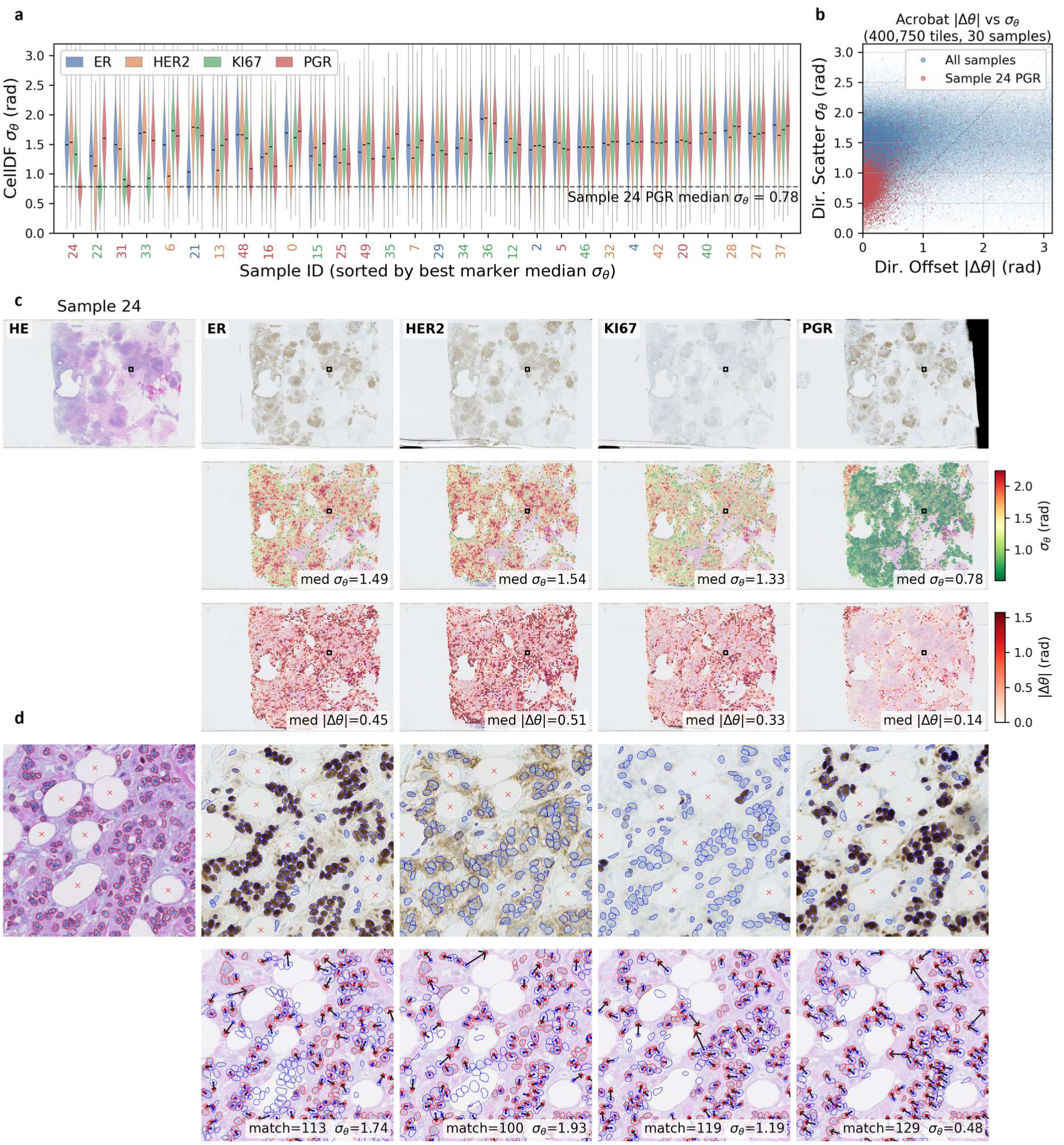
Serial-section CellDF matching quality across 30 Acrobat samples with four IHC markers. **(a)** Per-tile *σ*_θ_ violin plots for 30 samples × 4 markers (ER, HER2, KI67, PGR), sorted by the best marker’s median *σ*_θ_. Each sample position shows four violins (one per marker); x-axis labels are colored by the best-performing marker. Black dashes indicate per-marker medians. Dashed horizontal line: best sample–marker median *σ*_θ_ (Sample 24, PGR, 0.78 rad). **(b)** Per-tile |*Δθ*| versus *σ*_θ_ scatter pooled across all 120 pairs; red points highlight tiles from Sample 24 PGR. **(c)** Sample 24 WSI-level heatmaps across all four markers (columns: HE, ER, HER2, KI67, PGR). Row 1: WSI thumbnails with tile location box. Row 2: WSI *σ*_θ_ heatmaps with per-marker median *σ*_θ_ annotated. Row 3: WSI |*Δθ*| heatmaps with per-marker median |*Δθ*| annotated. **(d)** Sample 24 tile-level zoom-in across all four markers. Row 1: tile patches with cell contours (red: target, blue: warped); red crosses indicate manually annotated landmarks. Row 2: HE patches with target (red) and warped (blue) contours and CellDF match arrows (black), with per-tile *σ*_θ_ and match count annotated.

The four markers behaved heterogeneously across samples. In some samples, one marker yielded a distinctly lower median *σ*_θ_ than the other three; in others, all four markers produced comparably elevated values. Because Acrobat does not provide the serial-section order, we cannot know which (if any) of the four IHC markers corresponds to the section physically closest to HE; such determinations therefore require case-by-case inspection. We therefore examined Sample 24 in detail, the sample with the overall lowest median *σ*_θ_ (PGR) among all 120 pairs, where the other three markers showed substantially higher values.

Per-marker WSI heatmaps of *σ*_θ_ and |*Δθ*| showed that PGR exhibited consistently lower values across the tissue than ER, HER2, and KI67 (Fig. 4c). Zooming into representative tiles (Fig. 4d) revealed different underlying causes. In the ER and HER2 sections, the local cell composition differed from the HE section, most visibly in the number and distribution of adipocytes, indicating tissue-level differences between sections rather than purely non-rigid registration error. KI67 was intermediate, with cell composition closer to HE but still elevated *σ*_θ_ across much of the WSI. PGR alone matched HE in both cell composition and displacement consistency, producing arrows comparable to HyReCo same-section tiles. This pattern was also observed in Samples 22 and 31.

### 4.6 Matching Quality Control: Two-Stage Filtering

Even when CellDF identifies the plausibly closest IHC section to HE within a multi-marker dataset, the two sections are not physically identical, and localized tissue-level differences inevitably produce incorrect matches. Filtering these unreliable matches is therefore required before IHC labels are transferred downstream. We constructed a two-stage MAD-based filter that removes statistical outliers at the tile level within a WSI and at the match level within each tile (Fig. 5a).

**Fig. 5.**
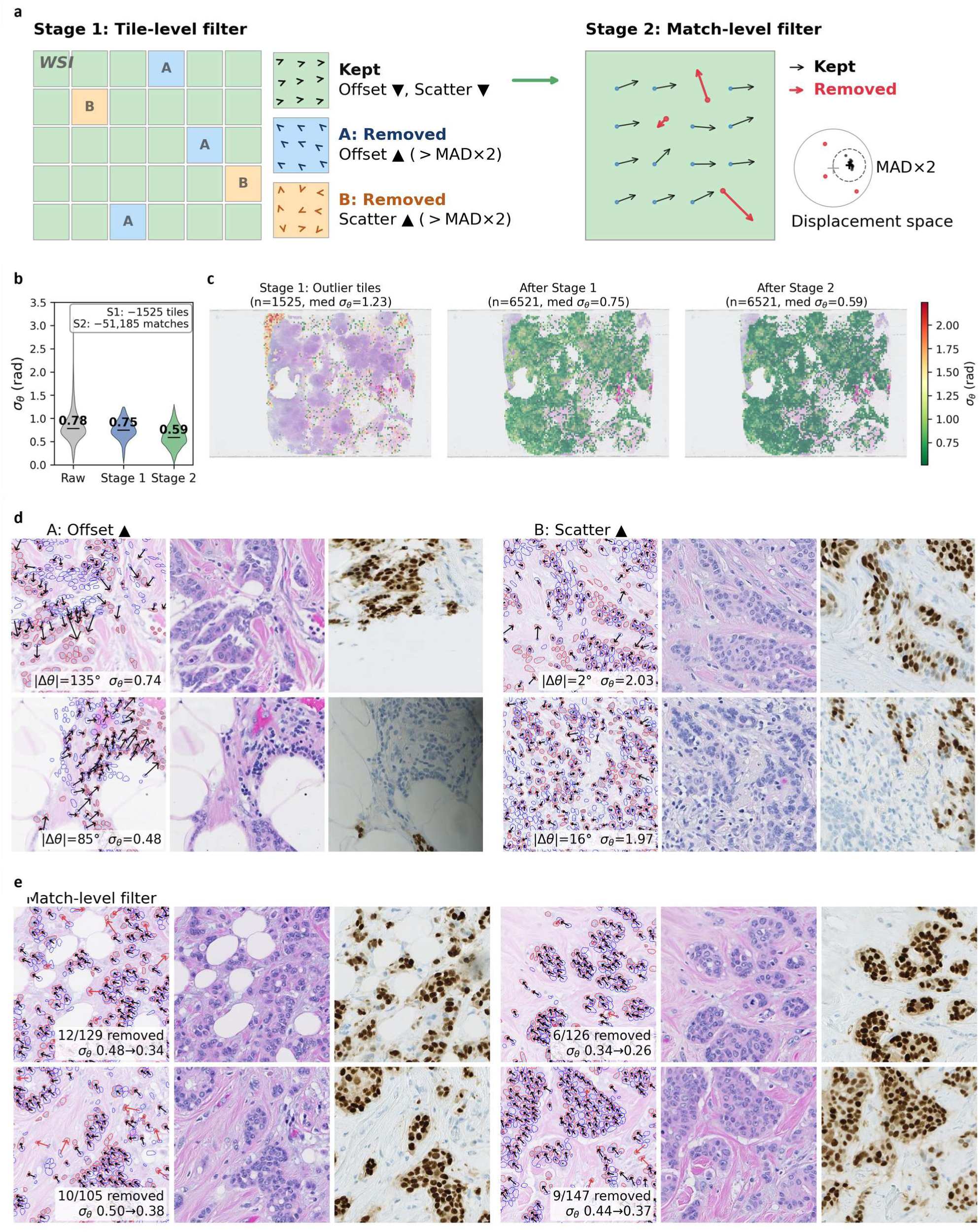
Two-stage MAD-based filter for matching quality control. **(a)** Schematic of the filter. Stage 1 (tile-level): a grid of tiles illustrates the WSI-scale screen, where each tile is classified as kept (green, |*Δθ*| low and *σ*_θ_ low), criterion-A outlier (blue, |*Δθ*| > MAD × 2), or criterion-B outlier (orange, *σ*_θ_ > MAD × 2). Three legend tiles with sample displacement arrows illustrate the three conditions. Stage 2 (match-level): a single kept tile zoom-in shows kept matches (black arrows) and outlier matches (red arrows) whose displacement residuals exceed MAD × 2; the adjacent displacement-space inset plots all matches as points relative to the tile median, with the dashed circle marking the MAD × 2 threshold and the cross marking the origin. **(b)** Per-tile *σ*_θ_ violin distribution for Sample 24 PGR at three stages: raw, after Stage 1, and after Stage 2. Medians and tile counts are annotated. **(c)** WSI *σ*_θ_ heatmaps (colormap shared with Fig. 4): (left) outlier tiles removed at Stage 1, (middle) surviving tiles after Stage 1, and (right) surviving tiles after Stage 2. Per-panel tile count and median *σ*_θ_ are annotated. **(d)** Representative tile-level outliers from Stage 1: left block (A: |*Δθ*| high) and right block (B: *σ*_θ_ high), each showing two tiles with target HE, warped IHC, and HE overlay with cell contours (red: target, blue: warped) and CellDF match arrows (black). **(e)** Representative tiles from Stage 2 match-level filtering (four tiles). Each tile shows, from left to right, HE overlay with kept (black) and removed (red) match arrows, Target HE, and Warped IHC. Per-tile counts of removed matches and the change in *σ*_θ_ before and after Stage 2 are annotated on the overlay.

Applied to Sample 24 PGR (607,341 CellDF matches across 8,046 qualified tiles), the filter progressively reduced the median per-tile *σ*_θ_ from 0.78 in the raw data, to 0.75 after Stage 1 (1,525 outlier tiles removed), and to 0.59 after Stage 2 (51,185 matches removed from the 6,521 surviving tiles), a 24% reduction from the unfiltered baseline (Fig. 5b), with the *σ*_θ_ heatmaps becoming progressively more homogeneous across the tissue (Fig. 5c). Representative tile-level outliers displayed the two distinct failure modes targeted by Stage 1: criterion-A tiles exhibiting directional deviation from the global field, and criterion-B tiles with arrows pointing in random directions (Fig. 5d). Within the surviving tiles, Stage 2 removed sporadic outliers embedded in otherwise well-registered regions (Fig. 5e). Across both stages, 16% of the original matches were removed (607K → 510K), retaining the majority of correspondences for downstream IHC label transfer.

### 4.7 Label Transfer and MLP Classification: Downstream Utility

As a proof-of-concept demonstration of downstream utility, we performed IHC label transfer and then predicted the transferred labels from HE cell embeddings on Sample 24 PGR, the best-matched pair identified in the serial-section analysis.

Among 510K matches surviving the two-stage filter, 55.4% of warped IHC cells were labeled Positive (bbox–DAB overlap ≥10%), 40.0% Negative (0%), and 4.6% Ambiguous (0–10%, excluded from training). The 486,567 Positive/Negative cells were transferred to their CellDF-matched Target cells (Fig. 6a), leaving 349,000 unlabeled Target cells (including Ambiguous-matched and unmatched cells) as the prediction set. We trained an MLP on the CellViT++ embeddings of labeled Target cells to predict labels for the prediction set (Fig. 6b), the Uniform Manifold Approximation and Projection (UMAP) of the labeled embeddings indicating that the two classes are separable in embedding space (Fig. 6c).

**Fig. 6.**
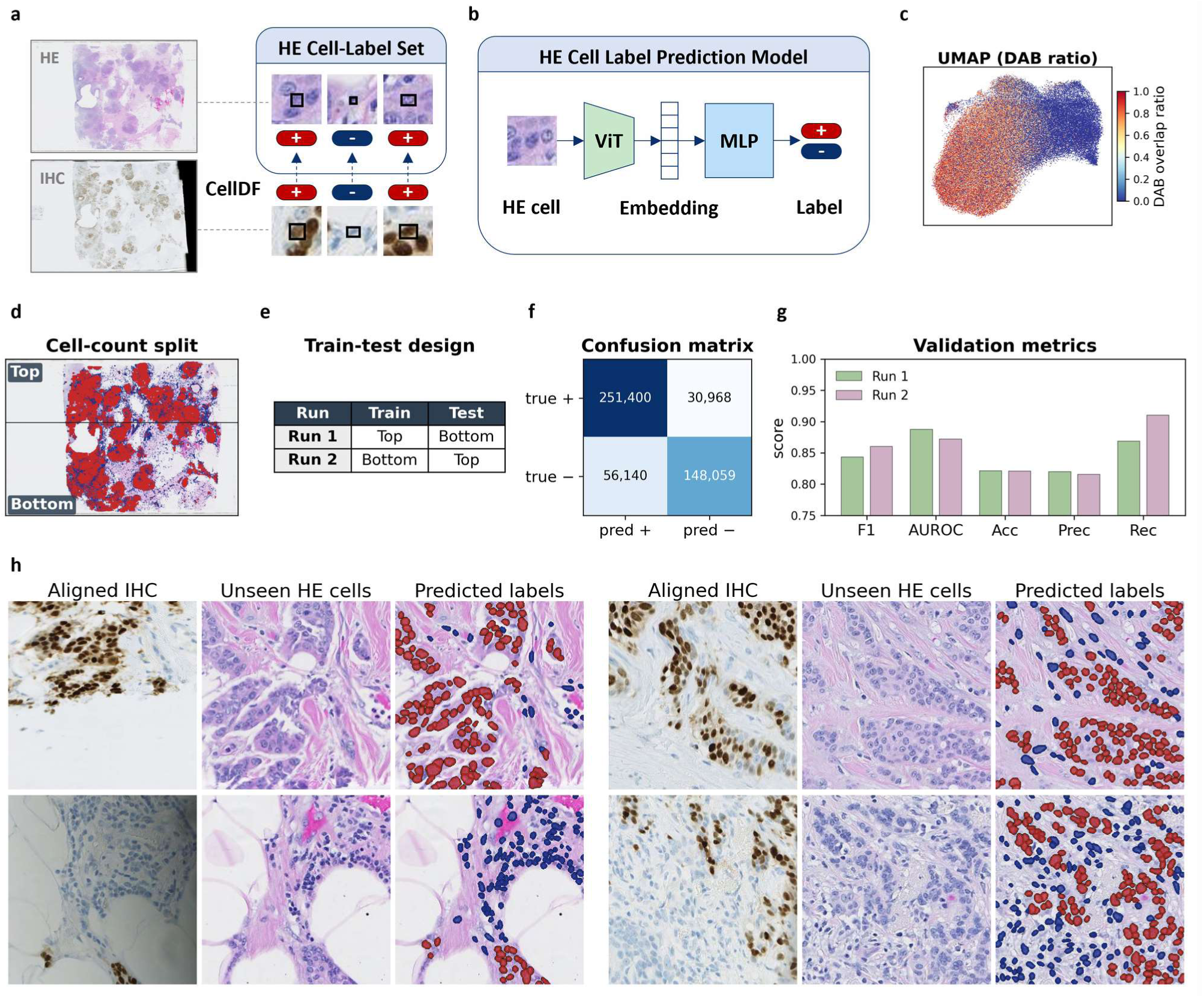
Per-cell IHC label transfer and MLP-based extension to filter-out regions. Sample 24 PGR throughout. **(a)** Cell-Label set construction. From the CellDF-matched filter-in pairs, each warped IHC cell is labeled IHC+ or IHC− using the nucleus bounding box, and the label is transferred to its matched HE target cell. **(b)** Label prediction model. A 1280-dim HE-cell embedding extracted by the SAM-H ViT is fed to a multi-layer perceptron (MLP) for binary IHC prediction. **(c)** UMAP projection of HE-cell embeddings (n = 100,000, stratified subsample) colored by IHC overlap ratio. **(d)** Cell-count split. The HE WSI is split horizontally at the y-median of labeled cells, giving Top and Bottom partitions with equal cell counts, with labeled cells overlaid. **(e)** Train–test design. Two MLPs are trained: Run 1 on Top and tested on Bottom; Run 2 on Bottom and tested on Top. **(f)** Confusion matrix summed over Run 1 and Run 2. **(g)** Validation metrics per run: F1, AUROC, accuracy, precision, recall. **(h)** MLP predictions on tiles not used in training (Stage-1 outlier tiles). An MLP trained on the full filter-in set is applied to all HE cells; four example outlier tiles are shown.

To assess within-sample generalization, we applied a cell-count split: the WSI was partitioned at the y-median of labeled cells into Top and Bottom halves of equal labeled-cell count (Fig. 6d). Two MLPs were trained with Top/Bottom reversed as train and test (Run 1 and Run 2; Fig. 6e), achieving an average F1 of 0.85 and AUROC of 0.88 across the two runs (Fig. 6f, 7g). A separate MLP trained on all 486,567 labeled cells was then applied to predict labels for previously unseen cells (Fig. 6h).

We repeated the same procedure on three further sample-marker pairs from the Acrobat 30-sample set, chosen as those that were both sufficiently well-matched and positive for the marker: average half-cut F1 and AUROC were 0.85 and 0.89 for Sample 48 (PGR), 0.83 and 0.89 for Sample 21 (ER), and 0.75 and 0.85 for Sample 31 (KI67). In every case the transferred labels were recoverable from HE within the sample, a prerequisite for using the pipeline output as a cell-level labeling resource.

## 5. Discussion

### 5.1 Principal Findings

This study treats serial-section IHC, the largest available pool of HE-IHC pairs in pathology, as material that a laboratory can turn into cell-level labels on its own cases, contingent on solving cell-level matching at WSI scale and assessing its reliability without ground-truth correspondences. CellDF estimates a locally adaptive residual displacement field through iterated kernel regression on each HE cell’s *K* nearest IHC candidates, producing matches that are coherent at patch scale and tractable at WSI scale where pairwise methods are not. The within-tile distribution of the resulting displacements supplies two diagnostic statistics, the directional scatter *σ*_θ_ and the between-tile angular deviation |*Δθ*|, that localize matching quality more finely than landmark-based median TRE and motivate a two-stage MAD filter; on the worked Acrobat example this filter removed roughly one sixth of the raw matches while reducing the median per-tile *σ*_θ_ by 24%. Discarding this fraction is acceptable in this regime because the serial-section resource is abundant and the intended use is training data, for which label precision matters more than complete coverage. In multi-marker serial-section data with unrecorded section adjacency, the same *σ*_θ_ statistic identifies which marker, if any, lies physically near enough to HE to support cell-level transfer. As a downstream demonstration on a worked Acrobat sample-marker pair (Sample 24 PGR), the IHC labels transferred through this pipeline trained an MLP that generalized within the sample at F1 0.85 and AUROC 0.88 under a held-out cell-count split. Taken together, these results establish that the resulting pairs of HE cells and their IHC labels are consistent enough to function as a labeling source within the worked sample, rather than verified correct against a cell-level ground truth that serial sections do not provide.

### 5.2 CellDF in the Algorithmic Landscape

CellDF treats cell matching as the estimation of the residual displacement that remains after WSI registration. Because registration has already brought corresponding cells into the same micrometer-scale neighborhood, the residual is small and varies smoothly across tissue, and the matching problem reduces to estimating that smooth field rather than searching the whole slide for each pairing. At every query cell, the predicted displacement is a Gaussian-weighted average of the displacements observed among nearby matched cells, and the query selects, from a small set of nearby candidates in the other modality, the one whose displacement best agrees with the local prediction. The selection and the field estimate refine each other and are iterated until they settle.

This places CellDF in the non-parametric kernel-regression family of non-rigid registration (Nadaraya, 1964; Watson, 1964), alongside the deformable variant of Coherent Point Drift (Myronenko and Song, 2010). The deformable CPD and CellDF in fact estimate the same kind of smooth field; the difference is which displacements each averages over. CPD couples every source cell with every target through a global Gaussian mixture model, and this pairwise coupling is what becomes prohibitive at whole-slide cell counts. CellDF carries out the same Gaussian-weighted estimation but only over each query cell’s *K* nearest candidates, retaining the locally adaptive character of the family while bypassing the global coupling. This restriction is what makes the framework usable at WSI scale, and it relies on prior registration having already placed each true correspondent within the candidate set.

Both RANSAC (Fischler and Bolles, 1981) and CellDF set a robust global initial estimate: each pools the vectors that every cell forms to its *K* nearest candidates and reduces them to a single global translation, differing only in the estimator, inlier consensus for RANSAC and geometric median for CellDF. The RANSAC baseline keeps this global vector as the final prediction; CellDF instead iterates local refinement on top of it, so switching the refinement off reduces CellDF to that baseline. Without local adaptation, local deviations are absorbed into the single global model, suppressing *σ*_θ_ where matching is in fact unreliable, so a global model is unfit as the matcher when *σ*_θ_ is also the registration-quality readout, because the matcher partially masks the metric it is measured by.

Beyond the matcher and its scaling lies a difference of objective. An established way to reach whole-slide scale with a pairwise method such as CPD is to fit it over a subsampled set of correspondences and interpolate the dense field elsewhere. Such an approach targets whole-slide registration, for which a dense, smoothly interpolated field is the natural output and landmark TRE the appropriate measure. Cell-level labeling imposes a different requirement: where serial sections genuinely lack a corresponding cell, a smooth field still assigns a displacement, but a label read from it there refers to no actual cell. CellDF is built around this distinction, reading per-tile displacement consistency to withhold labels in such regions rather than emit them and reporting reliability as a locally resolved statistic. Trading full coverage for the ability to localize and abstain on unreliable regions is what separates a labeling objective from a registration objective.

### 5.3 Per-Tile Statistics as a Ground-Truth-Free Quality Readout

The motivation for introducing *σ*_θ_ was practical. On the HyReCo dataset, several slides with apparently acceptable median TRE produced visibly noisy matches in localized regions of restaining damage. The per-sample median *σ*_θ_ correlated only moderately with the per-sample median TRE (Spearman *ρ* = 0.38), and four of the six samples flagged as *σ*_θ_-elevated had low median TRE. Sample 359 is the cleanest illustration: a globally moderate median TRE coexisted with a localized tissue-damaged region in which *σ*_θ_ rose sharply, and the surrounding tiles were affected as a consequence. Because median TRE summarizes a sparse landmark set with a single number, it cannot in principle separate these failure modes; *σ*_θ_ does so by reading the dense displacement field that matching itself produces. The two metrics report on different things, the agreement of registration with a sparse landmark set and the local coherence of matched displacements across all detected cells, which is why they are complementary rather than redundant.

*σ*_θ_ alone has a corresponding blind spot. In a tile where a small number of coincidentally aligned matches survive in an otherwise poorly registered region, the within-tile scatter can remain low even though the tile’s median displacement disagrees with the rest of the slide; conversely, a tile may scatter widely yet have a median displacement aligned with the global direction. Reading both the between-tile angular deviation |*Δθ*| and the within-tile scatter *σ*_θ_ catches either case, and the tile-level filter applies both criteria for the same reason. A second match-level stage then trims residual outliers within tiles that the tile-level stages let through. On the Acrobat Sample 24 PGR example, the full two-stage filter removed 16% of raw matches while reducing median per-tile *σ*_θ_ by 24%, the operating use of the same statistics that drove the diagnosis.

### 5.4 Marker Selection in Multi-Marker Serial-Section Datasets

The Acrobat application showed that the same statistics behave in a qualitatively different regime when the two modalities come from physically distinct sections. Per-tile *σ*_θ_ shifted substantially upward (Acrobat medians 0.78 to 1.80 rad versus HyReCo Sample 281 at 0.40 rad), and the |*Δθ*| distribution broadened substantially as well. This shift was expected, since serial sections sample non-identical cell populations, but the variation across the four markers within a single sample is as large as the gap between same-section and serial data, so marker choice can matter as much as that difference. In Sample 24, the PGR section had a median *σ*_θ_ close to the HyReCo same-section range while the other three markers were substantially elevated. Because Acrobat does not record the physical order of the serial sections, we cannot independently verify that PGR was the section physically nearest to HE; the per-marker heatmap evidence is consistent with that interpretation but not a proof, and the same pattern appeared in Samples 22 and 31. This turns the same statistic that diagnoses local registration failure on a single pair into a concrete pre-screening criterion across pairs: in multi-marker serial-section datasets where adjacency is unknown, *σ*_θ_ identifies which marker, if any, is likely close enough to HE to support cell-level label transfer.

### 5.5 Label Transfer as Proof of Concept

The Sample 24 PGR analysis was scoped as a proof of concept. After two-stage filtering, 486,567 matched cells provided Positive or Negative IHC labels for an MLP trained on CellViT++ embeddings (Hörst et al., 2025); the cell-count split gave F1 0.85 and AUROC 0.88, and the corresponding full-WSI MLP produced predictions for the 349,000 unlabeled cells in the prediction set. Reading these numbers requires three caveats. First, the labels treated as ground truth in evaluation are themselves transferred from a serial section through the same CellDF pipeline, so they encode the joint quality of registration, matching, and DAB labeling rather than an independent biological standard (Frénay and Verleysen, 2014; Shi et al., 2024; Campanella et al., 2019). Second, cells were partitioned spatially into Top and Bottom halves rather than at random, so that adjacent cells with correlated embeddings do not straddle the train and test sets and inflate the score (Bussola et al., 2021); this within-sample partition controls for the absence of a held-out test sample but still does not address cross-sample generalization (Roberts et al., 2017). Third, ambiguous-overlap exclusion improves training-data purity but means the prediction set is enriched for cells whose true label is genuinely uncertain, so the MLP’s behavior on that set cannot be directly compared with its evaluation on the labeled half. The honest claim is that the pipeline output is consistent enough to train an MLP that generalizes within the sample, not that the labels are correct in an absolute sense.

### 5.6 Tuning HE-Only Models on Labels Generated in the Same Laboratory

CellDF outputs directly address what HE-only molecular and functional predictors generally lack: given a serial IHC section, its output is HE cells paired with measured IHC labels, with each pairing tagged by a match-reliability estimate from the same per-tile statistics that drove label transfer. Because the IHC section is cut and stained where the HE section is, this material is produced by the same laboratory that would use it, on the cases that laboratory actually sees. Several uses follow.

The most direct use is the classification head of a cell detector. Modern detectors such as CellViT++ (Hörst et al., 2025) pair a SAM-derived backbone (Kirillov et al., 2023) with a lightweight classification head trained on HE-only datasets such as PanNuke (Gamper et al., 2019), which currently cover morphological cell types. CellDF outputs extend this training material along a molecular-phenotype axis (positivity for ER, PGR, HER2, Ki-67), allowing the same linear-classifier framework to be trained for markers and case mixes that public HE-only datasets do not cover, from the laboratory’s own sections and without manual annotation. The HE cells with their match-reliability tags, together with the SAM embeddings already computed by the upstream detector, form a paired resource that is directly compatible with the classification heads of pathology foundation models (Chen et al., 2024; Vorontsov et al., 2024; Lu et al., 2024; Xu et al., 2024), and that accumulates across cohorts.

Virtual staining is a second area. Recent generative models read enough spatial context to tolerate moderate HE-IHC misalignment (Ma et al., 2026), yet severely misregistered pairs still enter the training set and work against that design. For this image-to-image setting, CellDF’s per-tile reliability estimate flags where HE and IHC correspond well at cell resolution, so training can retain the well-matched regions and exclude or down-weight the severely misaligned ones instead of learning from them. This lets virtual staining draw training data from the serial-section archives that clinical practice produces in abundance, rather than being limited by their misalignment.

A reasonable objection is that cell-level matching is valid only when the HE and IHC sections are sufficiently adjacent. We accept this premise and turn it into an operating principle. The same per-tile *σ*_θ_ readout that estimates adjacency can route each pair, and each region within a pair, to the granularity it actually supports: cell-resolved label transfer where *σ*_θ_ indicates near-adjacency, and coarser region- or tissue-level labeling where it does not, instead of discarding the less-adjacent material outright. A heterogeneous serial-section archive can thereby contribute to a downstream dataset at mixed granularity rather than only through its closest pairs.

### 5.7 Limitations

CellDF is built on two preconditions that lie outside the matcher itself: that prior WSI registration has brought each true correspondent within the *K*-nearest candidate set, and that cell detection produces consistent positions in both modalities. When both hold, the *K*-nearest restriction delivers a direct computational gain over pairwise methods and the displacement-consistency metrics supply a ground-truth-free quality readout; when either fails, the pipeline neither detects nor recovers the failure. In regions of tissue folding or extensive damage the registration assumption breaks down. The magnitude of the residual itself depends on registration resolution: the same-section proof of concept was registered at a downsampled (2048-pixel) scale, and finer registration would shrink the residual, possibly to where nearest-neighbor assignment suffices on same-section data. CellDF is not intended to compensate for coarse registration; its role is for serial sections, where the residual reflects non-identical cell populations that no registration resolution removes, and at whole-slide scale, where registration is necessarily downsampled while the matcher and its per-tile reliability operate at cell resolution. In regions where the cell detector misses cells in both modalities the tile drops out of *σ*_θ_ computation, and when detections persist but correspond to non-overlapping tissue the tile produces high *σ*_θ_ that cannot be distinguished from registration error. Matching quality itself was assessed through these displacement-consistency statistics rather than through ground-truth correspondences, because such correspondences are not available at the scale of the analysis; the metrics are validated against landmark-based median TRE on HyReCo and against visual inspection on a handful of regions. For Acrobat, the absence of recorded serial-section order means that the worked result cannot be directly contextualized against an adjacency truth; the marker-selection use of *σ*_θ_ is therefore framed as an operating principle rather than a benchmarked claim. Of the downstream uses outlined above, only the cell-typing-classifier route was demonstrated, and only as a within-sample proof of concept: a held-out cell-count split on the worked Sample 24 PGR pair and on three further sample-marker pairs, confirming that the transferred labels carry a consistent signal learnable within a sample. This validates the internal consistency of the labeling output rather than its value as accumulated training data, the intended use, which would require assembling CellDF outputs across many samples and evaluating a downstream model trained on them.

### 5.8 Future Directions

Several extensions to the matcher and to the metrics it produces follow naturally. An adaptive bandwidth that varies with local cell density rather than being held fixed across a slide could let regions with few detected cells gather enough nearby matches for a stable local estimate, while removing the need for per-dataset tuning (Brockmann et al., 1993). The two diagnostic statistics and the two-stage filter built on them are sample-by-sample at present; aggregating them across many samples could yield dataset-level quality reports and inform per-sample thresholds. For multi-marker serial-section datasets, *σ*_θ_ measured between pairs of IHC sections could infer which sections sit physically adjacent to each other, even when the original order is not recorded; labels could then be transferred along this chain of adjacencies, so that IHC markers not next to HE can still contribute labels through their neighbors.

## 6. Conclusion

The combination of a sub-quadratic non-rigid matcher with a ground-truth-free quality readout moves cell-level cross-modal analysis closer to a routine operation on the serial sections that pathology laboratories already cut, rather than a special-case capability requiring purpose-built datasets. A laboratory can therefore generate cell-resolved molecular labels on its own cases and tune HE-only models on them. Serial sections provide no cell-level ground truth against which those labels can be checked, so the quality readout determines which of them are usable.

## Declarations

### CRediT authorship contribution statement

Eunji Jang: Conceptualization, Methodology, Software, Validation, Formal analysis, Investigation, Data curation, Visualization, Writing – original draft, Writing – review & editing. Yong-Min Huh: Conceptualization, Supervision, Project administration, Funding acquisition, Writing – review & editing.

### Data availability

This study analysed only publicly available datasets and generated no new patient data. The HyReCo dataset of restained histological serial sections, described by Lotz et al. (2023), is deposited at IEEE DataPort under a CC BY-SA 4.0 license and is available at https://doi.org/10.21227/pzj5-bs61; the subset used here is HyReCo-Additional, comprising 54 pairs of HE and PHH3 restained sections. The ACROBAT dataset of multi-stain breast cancer whole slide images, described by Weitz et al. (2024), is deposited with the Swedish National Data Service under a CC BY 4.0 license and is available at https://doi.org/10.48723/w728-p041. The CellDF implementation is publicly available under an MIT license at https://github.com/ejang2102/celldf, and the released version used for this work is archived at https://doi.org/10.5281/zenodo.22106732.

### Ethics statement

This work is a secondary computational analysis of publicly available, de-identified whole slide image datasets (HyReCo (Lotz et al., 2023) and ACROBAT (Weitz et al., 2024)). No new human participants, samples, or identifiable data were involved. Ethical approval for the original tissue collection was the responsibility of the respective dataset providers; no additional ethical approval was required for this study.

### Declaration of generative AI and AI-assisted technologies in the manuscript preparation process

During the preparation of this work the authors used Claude (Anthropic) to assist with drafting and editing the text of the manuscript. After using this tool, the authors reviewed and edited the content as needed and take full responsibility for the content of the publication.

## 7. References

Author-year (Harvard) style, which both MedIA and CMIG use. Inline citations have been converted from the former [shortform-key] placeholders to (Author et al., Year) form (2026-06-12); the Cited references below are sorted alphabetically by first-author surname. Same-author same-year entries are disambiguated with a/b (Wodzinski et al., 2024a/2024b). Journal names were abbreviated per LTWA (ISO 4) on 2026-08-07, as CMIG requires; single-word titles (Nature, Ecography) take no abbreviation, and conference proceedings and arXiv-only entries are outside the rule. Exact Elsevier punctuation (Surname, I., year after authors) remains a light revision-stage pass; full source data per reference is in manuscript/related_works/*/{key}.md.

